# Correlation scan: identifying genomic regions that affect genetic correlations applied to fertility traits

**DOI:** 10.1101/2021.11.05.467409

**Authors:** Babatunde S. Olasege, Laercio R. Porto-Neto, Muhammad S. Tahir, Gabriela C. Gouveia, Angela Cánovas, Ben J. Hayes, Marina R. S. Fortes

## Abstract

Reproductive traits are often genetically correlated. Yet, we don’t fully understand the complexities, synergism, or trade-offs between male and female fertility. Here, we introduce correlation scan, a novel framework for identifying the drivers or antagonizers of the genetic correlation between male and female fertility traits across the bovine genome. The identification of these regions facilitates the understanding of the complexity of these traits. Although the methodology was applied to cattle phenotypes, using high-density SNP genotypes, the general framework developed can be applied to any species or traits, and it can easily accommodate genome sequence data.

## Background

In animal genetics, insight into the nature of the genetic relationships between quantitative traits are important because they improve our understanding of complex traits and diseases [1, 2]. These relationships termed genetic correlations manifest when there is shared genetic influence between traits (i.e., pleiotropy) [3, 4] or when there is non-random association between loci (i.e., linkage disequilibrium (LD)) [5, 6]. Estimated genetic correlations provide information on how genome-wide genetic effects align between two complex traits [7]. Understanding the interplay between the genomic variants and their effects on quantitative traits can yield insights to improve the prediction of genetic merit and the understanding of complex traits’ biology [8-10]. Estimated genetic correlations have informed animal and crop breeding for many decades. For example, scrotal circumference is used as an indicator trait in beef cattle breeding because it is genetically correlated with female fertility traits [11]. Nevertheless, we still have a limited information of the regions across genome regulating the intersexual correlations between male and female fertility traits. Investigating these regions and leveraging on the resulting biological information could inspire new approaches in livestock breeding [12, 13].

Over the past 100 years, different methods have been employed to estimate the genetic correlation between traits [14-17]. Traditionally, these correlations are estimated from pedigree data. However, genome-wide single nucleotide polymorphisms (SNPs) are often used in recent times [18]. It is possible to estimate across-sex correlation between traits and this research niche continues to attract interest among quantitative geneticists [19-21]. The resulting estimates from both within and across-sex analyses range from -1 to +1, indicating the strength and magnitude of the correlation between traits [22]. Despite more than a century of research on estimating this parameter, it is only very recently that studies attempt to identify the region(s) in the genome that underpin genetic correlations between traits [23-25].

In theory, we propose that various genomic regions will contribute to the overall genetic correlation between complex traits. Further, some regions will be driving the genetic correlation while others might antagonize it. For instance, if the genetic correlation between two traits is 0.70, some regions will yield a significant and positive correlation, say 0.90, while other regions may antagonize the overall estimate, and in that region the correlation could be -0.50. Also, some genomic regions may be neutral, say 0.02 and not significant for the correlation between the studied traits. Identifying driver and antagonizing regions are of particular interest if they are for two important traits which are unfavourably correlated, for example milk yield and fertility in dairy cattle. Identification of such regions could lead to more targeted genomic selection and rapid genetic gains for both traits. Current genomic tools have created a great opportunity to advance our knowledge of genetic correlations between complex traits, by investigating the regions in the genome that drive or antagonize these correlations.

Here, we introduce a framework termed “correlation scan”, which uses a sliding window methodology to uncover the genomic regions driving and antagonizing genetic correlations in beef cattle. We applied the method to male and female fertility traits and showcase how the outcomes of this methodology can be interpreted in downstream analyses to gain further insight about the studied traits and their relationships. Reproductive traits are often genetically correlated, and yet we don’t fully understand the complexities, synergism, or trade-offs between male and female fertility. To demonstrate the method, we used two pairs of reproductive traits with strong genetic correlations in two independent cattle populations from our previous study [26]. These traits are age at first *corpus luteum* (AGECL, i.e., female puberty) and serum levels of insulin growth hormone (IGF1 measured in bulls, IGF1b, or cows, IGF1c). These pairs of traits serve as example of a positive and a negative correlation between phenotypes measured in males and females, during pubertal development. The populations used in the study are formed by either Brahman (BB) cattle or Tropical Composite (TC) cattle, as described in our previous study [26].

## Results

### The total number of windows generated and analysed

Using the framework developed in our study (see Materials and Methods), genomic windows with their corresponding correlation estimates (**r**) for each pairwise trait in two beef cattle populations were identified. The total number of windows generated for all pairwise traits in BB was 5,558 and the number in TC was 6,876. For all windows, the chromosome coordinates and the corresponding **r** estimates in each of the two populations are presented in Additional file 1 (Table S1-S2). The **r** estimates for all windows were plotted against their genomic position (i.e., midpoint between the start and end position of each window) (Figure 1). Results are presented separately per cattle population and for each pair of traits investigated

**Figure 1.**
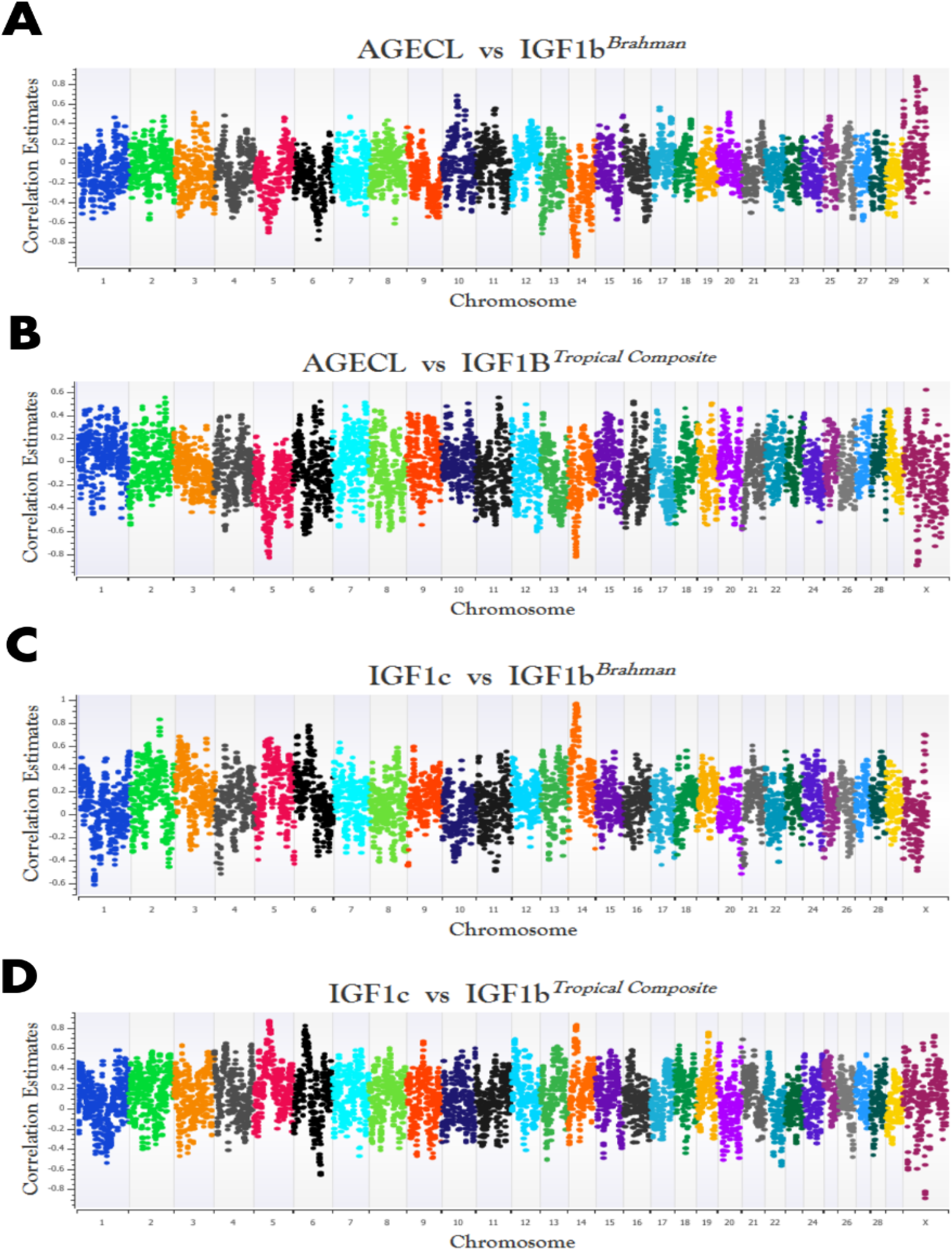
Genome plot of the regions driving and antagonizing trait correlations in Brahman (BB) and Tropical Composite (TC) for the pairwise traits (BB-AGECL vs IGF1b; **A**, TC-AGECL vs IGF1b; **B**, BB-IGF1c vs IGF1b; **C**, TC-IGF1c vs IGF1b; **D**). **AGECL**, age at first corpus; **IGF1**, serum levels of insulin growth hormone (measured in bulls, **IGF1b**, or cows, **IGF1c**). The correlation estimates were plotted on the y-axis and the genomic position (i.e., midpoint between the start and end position of each window) of each chromosome on the x-axis, according to the ARS_UCD1.2 bovine reference genome.

### Driver, antagonizing and neutral genomic windows affecting genetic correlations between fertility traits: permutation test

In order to identify drivers, antagonizing and neutral regions across the bovine genome, we performed permutation test by randomly reshuffling the Single Nucleotide Polymorphisms (SNPs) effects in each window across all chromosomes in 1,000 iterations for each trait. Then, we applied our framework on the randomized SNP effects and observed the **r** estimate across each iteration for each window. In most cases, the maximum and the minimum **r** estimates for each window at each iteration (i.e., rand 1 to rand 1,000) range between ±0.20. Therefore, we considered neutral windows with no significant effect on the trait correlation as windows with -0.20≤ **r** ≤0.20 estimates. The genomic plots of the **r** estimates resulting from the permutation test (rand 500 only) in each population are presented in Additional file 2 (Fig. S1). Additional file 3 (Table S3-S6) shows the numbers of windows, their chromosome coordinates, and **r** estimates as well as the maximum and minimum r estimates across the 1,000 iterations for each pair of traits in each of the two populations.

As a result of the permutation test, we considered significant windows with **r** estimates >0.2 and **r** estimates <-0.2. These thresholds were used to define the significant windows or regions (i.e., driver and antagonizing) from non-significant (i.e., neutral) windows or regions. Depending on the overall genetic correlation between traits, driver and antagonizing windows can be deduced: in driver windows, the **r** estimate has the same direction, positive or negative, as the overall genetic correlation; in antagonizing windows it is the opposite.

The number of significant driver windows for the correlation between AGECL and IGF1b was 1,636 in BB and 1,914 in TC cattle. The number of significant windows for the antagonizing was 547 in BB and 898 in TC cattle, for AGECL vs IGF1b. For the correlation between IGF1c and IGF1b, the number of significant driver windows was 1,931 in BB and 2,549 in TC cattle. The antagonizing windows was 402 in BB and 587 in TC cattle (IGF1c vs IGF1b). The numbers of neutral windows were as follows: 3,375 in BB, and 4,064 in TC for AGECL vs IGF1b; and 3,225 in BB, and 3,740 in TC for IGF1c vs IGF1b. See Table 1 for details on numbers of windows in the two beef cattle populations. In addition, the lists of windows with their chromosomal coordinates for all driver, antagonizing, and neutral regions are presented in Additional file 4 (Table S7-S18).

**Table 1:** The number of windows generated for the driver, antagonizing and neutral windows for each pairwise trait in Brahman and Tropical Composite population.

| Pairwise Trait | Number of windows (percentage to the total number of window) |  |  | Total number of windows |
| --- | --- | --- | --- | --- |
|  | Driver (%) | Antagonizing (%) | Neutral (%) |  |
| Brahman |  |  |  |  |
| AGECL vs IGF1b | 1,636 (29.40%) | 547 (9.84%) | 3,375 (60.72%) | 5,558 |
| IGF1c vs IGF1b | 1,931 (34.74%) | 402 (7.23%) | 3,225 (58.03%) | 5,558 |
| Tropical Composite |  |  |  |  |
| AGECL vs IGF1b | 1914 (27.84%) | 898 (13.06%) | 4064 (59.10%) | 6,876 |
| IGF1c vs IGF1b | 2549 (37.73%) | 587 (8.54%) | 3740 (54.39%) | 6,876 |

**AGECL**, age at first *corpus*; **IGF1c**, serum levels of insulin growth hormone measured in cow; **IGF1b**, serum levels of insulin growth hormone measured in bulls

For the correlation between AGECL and IGF1b (overall genome-wide correlation of -0.65 (BB) and -0.55 (TC), see Table 2), the largest **r** estimate for the driver windows was -0.96 (bovine chromosome (BTA)14: 23.04 - 25.29Mb) in BB and -0.91 (BTAX: 39.76 - 42.86Mb) in TC. For the antagonizing windows, the largest **r** estimate was 0.87 (BTAX: 40.87 - 43.88Mb) in BB and 0.61 (BTAX: 66.62 - 69.622Mb) in TC.

**Table 2:** Genomic correlations estimates and their corresponding standard error (s.e), number of animals and number of SNPs estimates.

| Pairwise traits | No of animals | Number of SNP | Genetic correlation (s.e) |
| --- | --- | --- | --- |
| *Brahman* |  |  |  |
| AGECL vs IGF1b | AGECL- 980 | 554K | -0.65 (0.13) |
|  | IGF1b- 964 |  |  |
| IGF1c vs IGF1b | IGF1c- 995 | 554K | 0.86 (0.11) |
|  | IGF1b- 964 |  |  |
| *Tropical Composite* |  |  |  |
| AGECL vs IGF1b | AGECL-996 | 686K | -0.55 (0.14) |
|  | IGF1b- 998 |  |  |
| IGF1b vs IGF1b | IGF1c- 1015 | 686K | 0.93 (0.11) |
|  | IGF1b- 998 |  |  |

AGECL, age at first *corpus*; IGF1c, serum levels of insulin growth hormone measured in cow; IGF1b, serum levels of insulin growth hormone measured in bulls; SNP, Single Nucleotide Polymorphisms; s.e, standard error

For the correlation between IGF1c and IGF1b (overall genome-wide correlation of 0.86 (BB) and 0.93 (TC), see Table 2), the largest estimate for the driver windows was 0.97 (BTA14: 22.68 - 24.96Mb) in BB and 0.87 (BTA5: 46.13-47.89Mb) in TC, while the estimate for the antagonizing was -0.62 (BTA1: 49.01 - 51.67Mb) in BB and -0.90 (BTAX: 65.64 - 68.39Mb) in TC. All **r** estimates are plotted in Figure 1.

### Genes and quantitative trait loci (QTL) within driver and antagonizing regions across the two populations

Defining driver and antagonizing regions separately for each pair of traits, allowed us to identify the genes and QTLs within these regions for each of the two beef cattle populations. The percentage of the overlapping genes (Figure 2) and QTLs (Figure 3) across both populations was studied. The percentages of genes shared across the significant regions in BB and TC were calculated as a function of the total number of genes in BB or TC, respectively, and so they differ (Figure 2 and 3).

**Figure 2.**
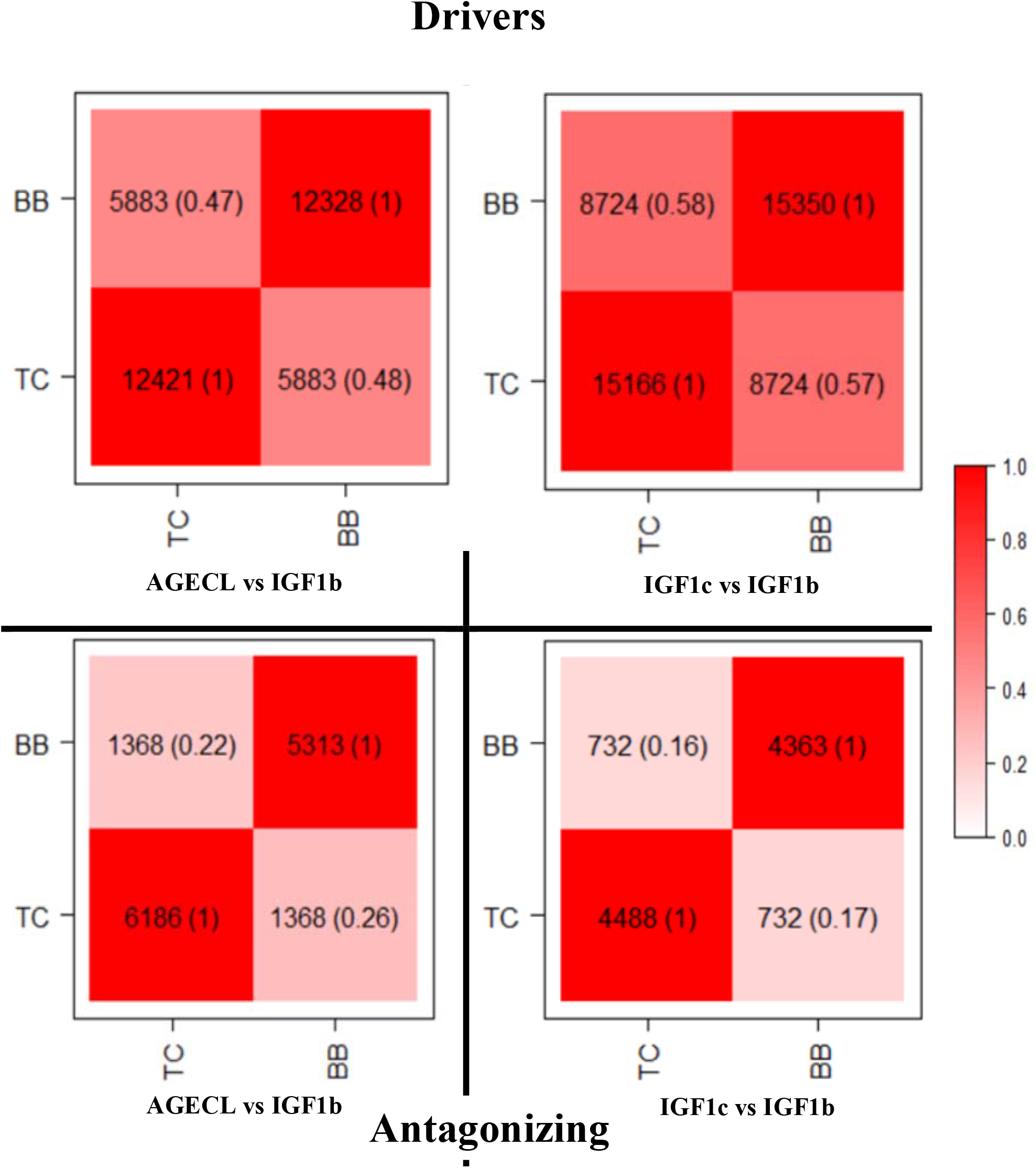
Genes annotated in the significant (i.e., driver and antagonizing) genomic regions identified as explaining the genetic correlations between male and female fertility traits in Brahman (BB) and Tropical Composite (TC) population. The overlaps between the two studied populations are in the diagonal of each plot for each pair of traits within the driver (above) and antagonizing (below) regions. The darker the colour within the squares, the higher the percentage of shared genes or QTLs. **AGECL**, age at first corpus; **IGF1**, serum levels of insulin growth hormone (measured in bulls, **IGF1b**, or cows, **IGF1c**).

**Figure 3.**
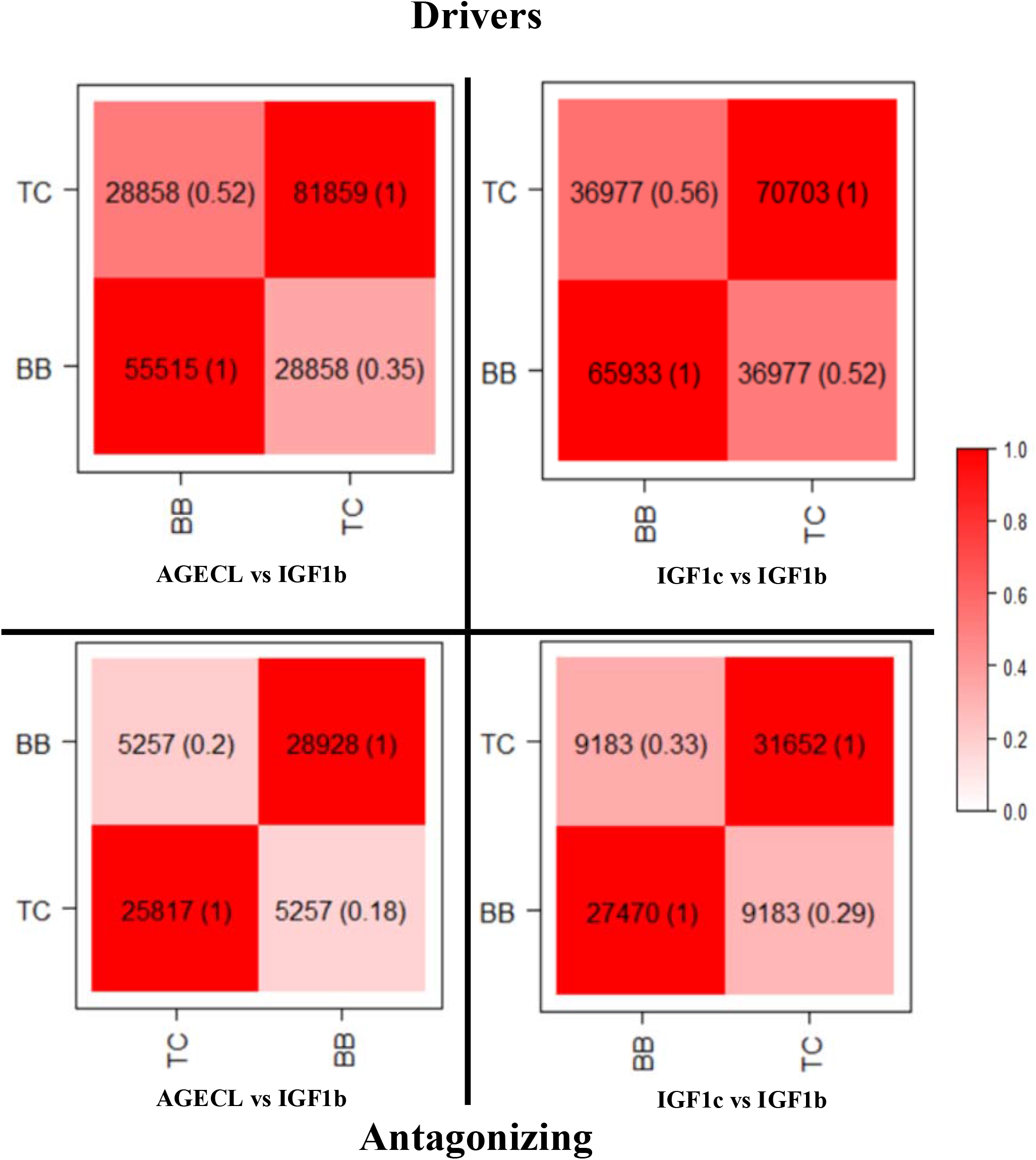
QTLs annotated in the significant (i.e., driver and antagonizing) genomic regions identified as explaining the genetic correlations between male and female fertility traits in Brahman (BB) and Tropical Composite (TC) population. The overlaps between the two studied populations are in the diagonal of each plot for each pair of traits within the driver (above) and antagonizing (below) regions. The darker the colour within the squares, the higher the percentage of shared genes or QTLs. **AGECL**, age at first corpus luteum; **IGF1**, serum levels of insulin growth hormone (measured in bulls, **IGF1b**, or cows, **IGF1c**).

The percentage of overlapping genes for each pair of traits in the two populations were as follows: for AGECL vs IGF1b driver regions, about 48% of the total number of genes annotated were shared between the two population, whereas, for the antagonizing regions, 22% of the gene annotated in BB were present in the TC population, and 26% of the genes annotated in TC were present in BB; for IGF1c vs IGF1b, the two populations shared about 58% of total number of genes annotated for the driver regions and about 17% were shared for the antagonizing regions.

The percentage of overlapping QTLs for each pair of traits in BB and TC population were as follows: for AGECL vs IGF1b driver regions, 52% of the QTLs annotated in BB were present in TC and 35% of the QTLs annotated in TC were present in BB, whereas, for the antagonizing regions, 20% of the QTLs annotated in BB were present in TC and 18% of the QTLs annotated in TC were present in BB; for IGF1c vs IGF1b, 56% of the genes annotated in BB were present in TC, and 52% of the genes annotated in TC were present in BB, whereas, for the antagonizing regions, 29% of the genes annotated in BB were present in TC and 33% of the genes annotated in TC were present in BB population.

### Functional classification of QTLs within genomic regions that explain the genetic correlations between male and female fertility

To infer biological function and mine the existing literature, we examined the types of QTL (milk, reproduction, production, meat and carcass, health and exterior) present in the significant genomic regions identified above using GALLO [27]. The most frequent QTLs across all pairwise traits in the two populations for the driver and antagonizing regions were QTLs related to milk production, accounting for about 30-51% in most cases. This was followed by reproductive QTLs accounting for about 13-48% and production QTLs comprising 6-24%. Other QTL types (Exterior, health and meat and carcass) accounted for a relatively small proportion of QTLs in the significant regions (Figure 4 and 5). In addition, we report the top 10 results for QTLs related to reproductive traits as these are relevant to our studied traits (Figure 4 and 5). Among these reproductive QTLs, traits related to puberty (i.e., age at puberty, scrotal circumference) were prevalent in both populations.

**Figure 4.**
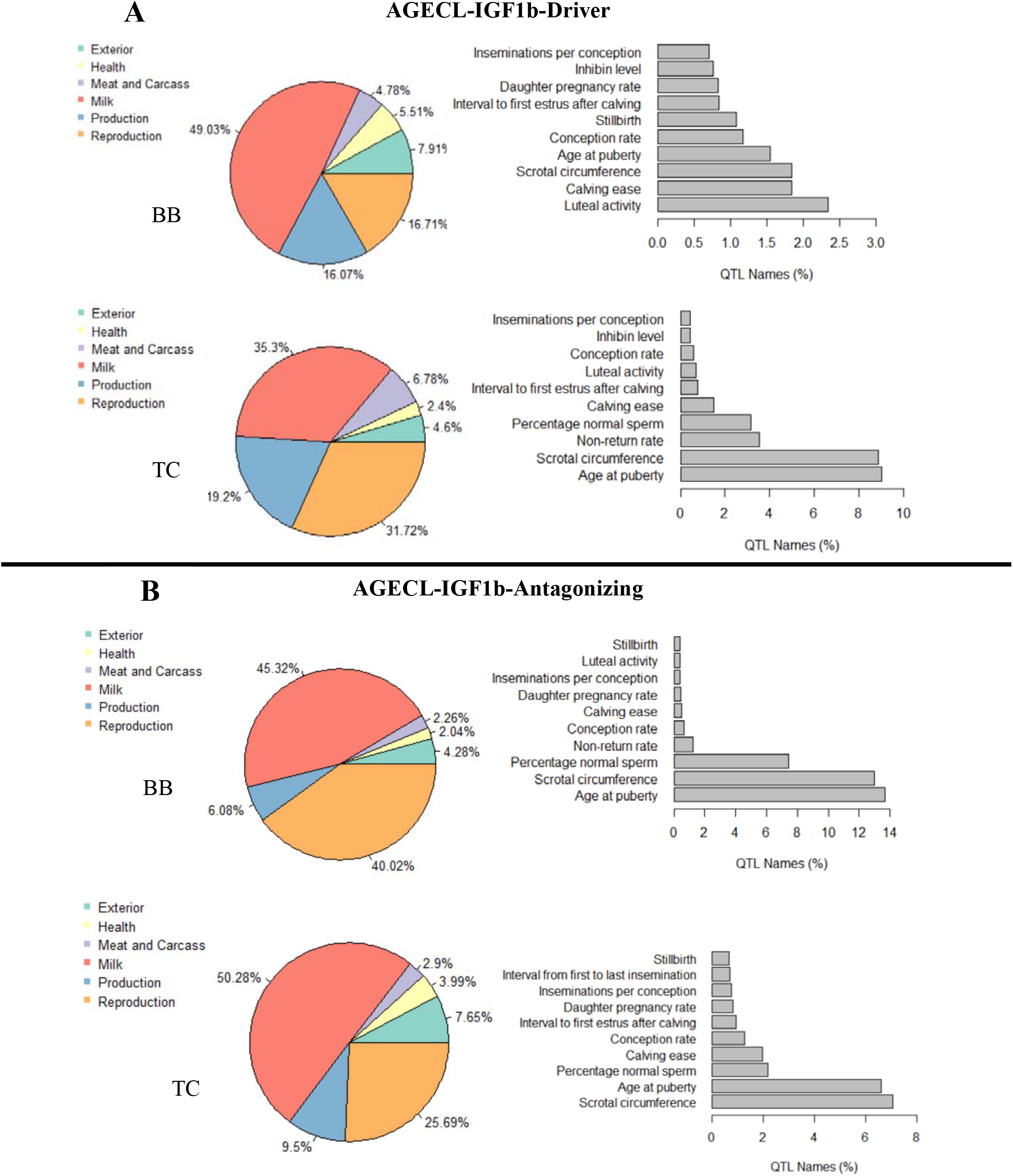
Percentage of QTL type (pie chart) and trait related to reproduction QTLs (barplots) for the QTL annotation results obtained for (A) AGECL vs IGF1b - driver, (B) AGECL vs IGF1b-antagonizing in Brahman (BB) and Tropical Composite (TC) population. **AGECL**, age at first corpus luteum, **IGF1**, serum levels of insulin growth hormone (measured in bulls, **IGF1b**, or cows, **IGF1c**)

**Figure 5.**
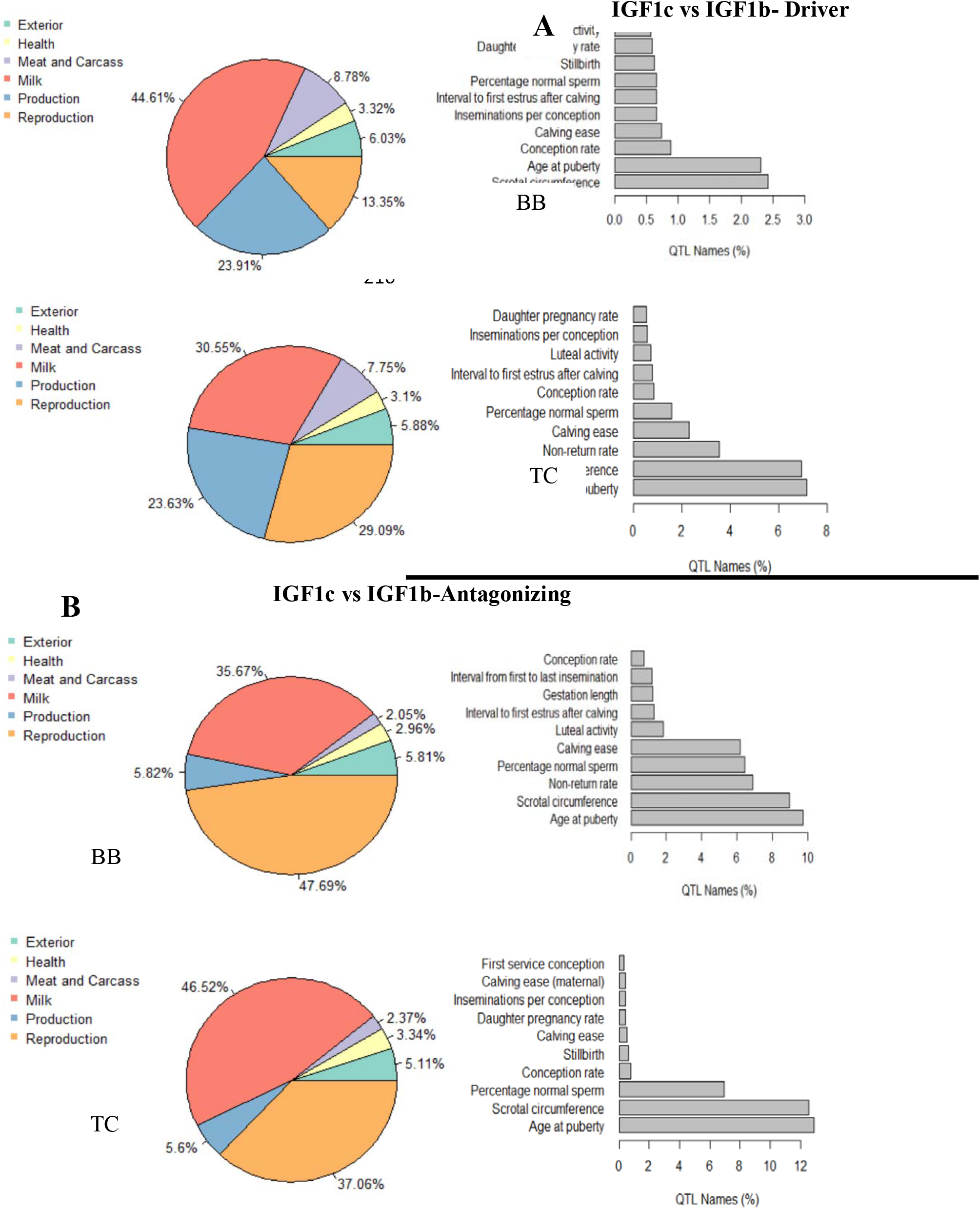
Percentage of QTL type (pie chart) and trait related to reproduction QTLs (barplots) for the QTL annotation results obtained for **(A)** IGF1c vs IGF1b-driver, (B) IGF1c vs IGF1b-antagonizing in Brahman (BB) and Tropical Composite (TC). **AGECL**, age at first corpus luteum, **IGF1**, serum levels of insulin growth hormone (in bulls, **IGF1b**, or cows, **IGF1c**).

### QTL enrichment analysis

We performed a chromosome-wide QTL enrichment analysis to further test the significance of the QTLs identified for all the driver and antagonizing regions in each cattle population, for each trait pair using GALLO [27]. Enriched QTLs for the studied traits span across most QTL types, indicating the presence of complex genetic mechanisms. The results of the chromosome-wide QTLs enrichment (FDR-corrected p-value≤0.05) for the driver and antagonizing regions for all pairwise traits in each population are presented in Additional file 5 (Table S19-26).

For the driver regions, the number of QTLs enriched over a wide range of chromosomes for AGECL vs IGF1b were 233 and 144 in BB and TC beef cattle population, respectively. The number was 227 (BB) and 220 (TC) for IGF1c vs IGF1b. For AGECL vs IGF1b, the most enriched chromosome (no of enriched QTLs in parenthesis) was BTA5 (36) and BTA14 (18) in BB and TC, respectively. IGF1c vs IGF1b also followed similar pattern with the result above, with BTA5 (41) being the most enriched chromosome in BB and BTA14 (51) as the most in TC.

For the antagonizing regions, the number of QTLs enriched across the bovine chromosomes for AGECL vs IGF1b were 127 and 178 in BB and TC beef cattle population, respectively. The number was 179 (BB) and 195 (TC) for IGF1c vs IGF1b. For AGECL vs IGF1b, the most enriched chromosome was BTA17 (14) and BTA26 (21) in BB and TC, respectively. For IGF1c vs IGF1b, however, BTA14 (23) was the most enriched chromosome for these regions in BB, whereas, in TC, BTA14 (23) was the most enriched.

To identify the common results and shared biology between the driver and the antagonizing regions, we also investigated the overlaps of the QTL types associated with the studied trait (i.e., reproduction) in the two populations. The relationship between the top 10 enriched reproductive QTLs in BB and TC are presented in Figure 6. Irrespective of the trait pair, for the driver regions, the reproductive QTLs in BB in most cases overlap with those identified in TC. However, for the antagonizing regions, not all reproductive QTLs in BB were found in TC beef cattle population.

**Figure 6.**
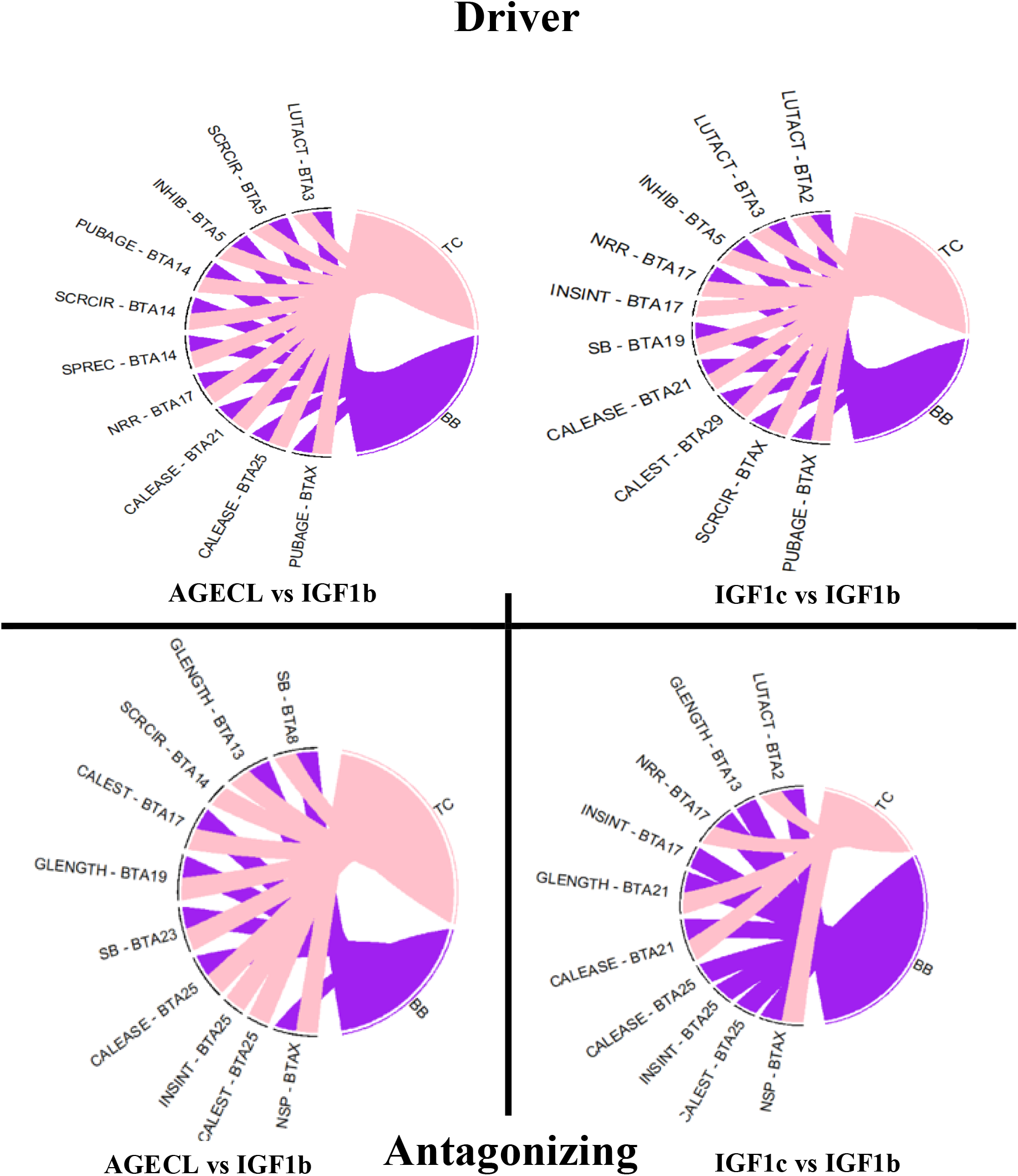
Chord plot showing the relationship between the top 10 enriched reproductive QTLs between Brahman (BB) and Tropical Composite (TC) for the driver (top) and the antagonizing (bottom) regions of the studied traits. **AGECL**, age at first corpus luteum, **IGF1c**, serum levels of insulin growth hormone measured in cow; IGF1b. LUTACT, Luteal activity; SCRCIR, Scrotal circumference; INHIB, Inhibin level; PUBAGE, Age at puberty; SPREC, Sexual precocity; NRR, Non-return rate; CALEASE, Calving ease; INSINT, Interval from first to last insemination; SB, Still birth; CALEST, Interval to first estrus after calving; GLENGTH, Gestation length, NSP, Percentage normal sperm.

### Functional enrichment analysis

Leveraging our methodology’s directionality of gene effects with Ingenuity Pathway Analysis (IPA; http://www.ingenuity.com), we identified the enriched canonical metabolic pathways enriched at Benjamini–Hochberg corrected p-values (BH-*P*-value) of p<0.01. The graphical presentation of the canonical metabolic pathways predicted by IPA to be enriched and the proportion of driver and antagonizing genes in each pathway for all pairwise traits investigated in each population are illustrated in Additional file 6 (Fig. S2-S9). Although IPA provided information about whether the predicted pathways were being activated or inhibited based on our data, we remain cautious when interpreting our results since the **r** estimates are not the same as gene expression values, and IPA was originally designed to mine gene expression data.

The number of pathways enriched for AGECL vs IGF1b was 49 in BB and 68 in TC. For IGF1c vs IGF1b, the number of enriched pathways was 156 and 87 in BB and TC, respectively. For AGECL vs IGF1b in BB, the top 5 enriched canonical metabolic pathways were cardiac hypertrophy signaling (Enhanced), toll-like Receptor signaling, IL-6 Signaling, hepatic fibrosis signaling pathway and STAT3 Pathway. In TC population, the top 5 enriched canonical metabolic pathways were breast cancer regulation by stathmin1, signaling by Rho family GTPases, opioid signaling pathway, endocannabinoid developing neuron pathway, and CREB signaling in neurons.

For IGF1c vs IGF1b in BB, the top 5 enriched canonical metabolic pathways were cardiac hypertrophy signaling, CREB signaling in neurons, thrombin signaling, estrogen receptor signaling, opioid signaling pathway and AMPK signaling. In TC, the top 5 enriched pathways were CREB signaling in neurons, cardiac hypertrophy signaling (enhanced), opioid signaling pathway, gαs signaling and breast cancer regulation by stathmin1. The enriched canonical metabolic pathways and all the genes involved in each pathway are available in Additional file 7 (Table S27-30).

## Discussion

Complex phenotypes, including fertility, consist of multiple genetically correlated rather than independent traits [6]. The interplay between traits involve many genomic regions, usually in a large and polygenic regulatory network [28-31]. Genomic signals that regulate (i.e., drive or antagonize) complex traits are widely spread across the genome, including near many genes without significant effect on the phenotype or disease [29]. In the present post-genomic era, unravelling the genomic regions that regulate complex traits and the metabolic pathways associated with these phenotypes has become an important aspect of genetic studies in humans and animals [32]. In this study, we developed a novel framework termed correlation scan to reveal the significant regions that either drives or antagonize the genetic correlations between traits, across the genome. In addition, this method can also reveal genomic regions with no effect on the studied traits (neutral windows). The framework developed uses best linear unbiased prediction (BLUP) solution of SNP effects to estimate the local correlations between studied traits. Local correlations are based on sliding windows of 500-SNPs. We applied these sliding windows approach to reproductive traits measured in two populations and subject the outcomes, significant windows, to further analyses using GALLO [27]. and IPA (http://www.ingenuity.com) to gain further insight about the biology of studied traits and their relationships. Although the methodology was applied to beef cattle traits, using high-density SNP chip genotypes, the general framework can be applied to any species, any traits, and it can easily accommodate sequence level data.

Our results agreed with the established notion that multiple loci regulate reproductive traits [33-35]. Also, the mode of action of these loci and the magnitude of their effect varies across the genome. While some regions had no effect on the genetic correlations under investigation, other loci drive or antagonize the relationships between male and female fertility. The identification of driver and antagonizing loci creates opportunities to further understand the molecular mechanisms affecting quantitative traits. For example, correlations estimated from SNP effects have allowed researchers to construct gene networks [36]. Thereby, these types of approaches contribute to linking genotype with phenotype.

The two beef cattle populations investigated in this study are distinct in terms of their genetic composition. Brahman (BB) cattle are typically of *Bos indicus* origin whereas TC beef cattle emanated from the crossing between Bos *indicus* and Bos *taurus* breed [37]. Despite these differences, we found that a considerable number of annotated genes and QTLs driving trait correlation overlaps across breeds, although with variations in the size of SNP effects. This corroborates the findings of Bolormaa, Pryce [38], where a substantial number of QTLs were found segregating in *Bos indicus* and composite cattle using the same dataset. In this present study, the top genomic signal driving trait correlation across all pairwise traits in BB were located on BTA14. The significant region contains a widely known and well-characterized QTL, including the *PLAG1* gene, reported to be associated with growth and reproductive traits in our populations and other studies [39-44]. In TC however, the top signal differs across traits and mostly spread across two or three chromosomes, although with considerable number of overlaps with BB. This could be partly due to the variations in the architecture of composite breed [45]. The genome of composite breeds usually contains new haplotypes emerging from generations of crossbreeding. Moreover, the contribution of the founder populations on chromosomes and specific genomic regions are usually unevenly distributed, which most likely shapes the genome of composite breeds [45]. In short, differences between BB and TC are likely to impact the results of our analyses. Breed differences are expected, and so when two breeds share a similar result, it enhances our confidence in calling significant windows for the interplay between male and female fertility traits.

Most genomic regions antagonizing the genetic correlations between male and female fertility traits were located on chromosome X. Gene expression on chromosome X differs across-sex, resulting in genomic sexual conflict [46-48]. Genes in these antagonizing regions include *PO1FB, ZNF711, APOOL, HDX, DACH2, FAM133A*, among others. These genes are associated with different disorders including infertility, reproductive deficiencies, primary ovarian failure [49-51]. When some of these genes are over-expressed, it can dysregulate the cristae morphology of the mammalian mitochondria [52]. Understanding how these antagonizing genes interact to influence (in)fertility could help improve the reproductive potentials of beef cattle.

In animal production, more research is carried out on milk production-related traits, thereby creating large proportion of records for these traits in the cattle QTL database. These volumes of records can create a bias in the QTLs representativeness [27]. The QTL enrichment analysis allows testing the significance of the QTL representative using chromosome-wide approach to detect specific genomic region with many QTLs for a specific trait. For example, taking the driver regions for AGECL vs IGF1b in BB, the top enriched QTLs was found in BTA5, harbouring 36 QTLs. These QTLs comprised 8 different QTLs for reproduction (inhibin level, scrotal circumference, interval of first estrus after calving, gestation length, insemination per conception, conception rate, daughter pregnancy rate, and pregnancy rate). These 8-traits listed here have been found to be correlated with puberty (studied traits) in cattle. For instance, inhibin is regarded as a biomarker for sexual development because it regulates spermatogenesis in both beef and dairy bulls [53, 54]. Moderate genetic correlation was found between inhibin and AGECL [55] and between inhibin and IGF1b [56] in BB.

Scrotal circumference has also been found to be a moderate predictor of AGECL and IGF1b in BB [21, 26]. Thus, BTA5 may be a candidate region for fertility in BB beef cattle population. Other enriched QTLs out of the 36 mentioned above include 8 different production traits (average daily gain, metabolic body weight, length of productive life, body weight, rump width, body depth, residual feed intake, and net merit). These traits are related to feed efficiency in cattle. Improving feed efficiency of beef cattle is a major concern for beef producers. A recent study from Canal, Fontes [57] found that heifers that efficiently utilize feed attain puberty early than less feed efficient ones. Moreover, heifers that attain puberty at a relatively younger age have the potential to conceive early in life and be more productive throughout their lifetime [58]. In addition, IGF1 is an effective selection tools to improve feed efficiency and other production related traits, allowing breeders to preselect animals that can utilize feed efficiently [59, 60]. Other enriched QTLs for BTA5 in BB are related to exterior (7), milk (6), milk and carcass (5) and health (3) traits. Of note, the objective of most beef cattle breeding programs is to change the genetic merit of their cattle for many traits of interest [61]. The recurrent association of the BTA5 with multiple traits could suggest complex genetic mechanisms such as pleiotropy, epistasis, hitchhiking effects, linkage disequilibrium etc., regulating this chromosomal region [62, 63]. Therefore, breeders could target BTA5 to select multiple traits without any antagonistic effect on other traits listed herein.

Another interesting result from this study is the shared biology between the two breeds relative to the traits under study. Despite breed differences, the enriched reproductive QTLs driving the genetic correlations between male and female fertility are the same for the two cattle populations (Figure 6). Most of the enriched QTLs are related to reproductive traits measured early life. A possible explanation could be that the reproductive phenotypes shared common fundamental biology in the two populations. For the antagonizing regions, however, most of the reproductive QTLs were breed specific depending on the trait pairwise. Perhaps, this could be partly explained by the diverse genetic composition of the two breeds. Understanding the genomic architectures driving these early-in-life male and female fertility traits and their known genomic antagonisms could foster effective selection for both traits in tropical breeds [64, 65].

The major challenge faced by researchers when analysing an overwhelmingly large amount of genomic data is how to extract meaningful mechanistic insights into the underlying biology characterizing the given trait under study. To increase the explanatory power of genomic studies, pathway analysis has become first choice, providing researcher with the ability to infer meanings to high-throughput genomic data [66]. Leveraging the directionality of gene effects from our method with IPA knowledge base, several biological pathways known to be involved in reproduction (i.e., studied trait) were significantly enriched for all pairwise traits investigated across the two breeds. These pathways include estrogen receptor signaling, p38 MAPK signaling, GnRH signaling, sperm motility, cAMP-mediated signaling, AMPK signaling, and androgen signaling. Although IPA provided information about the activation or inhibition state for the enriched canonical metabolic pathways with the use of the **r** estimates in place of the gene expression values, we are not sure if these pathways were being activated or inhibited since we don’t have information about the expression values of the genes in these pathways. For example, Rho GDP Dissociation Inhibitor (RHOGDI) pathway was the only significant signaling pathway found to be inhibited across breeds in all pairwise traits investigated using IPA comparison analysis. The RHOGDIs (RHOGDIα, RHOGDIβ and RHOGDIγ) are well-characterized as a negative regulator of Rho GTPases [67]. These Rho GTPases play pivotal roles within the cell, including cell migration, membrane trafficking, invasion, gene transcription, polarity, adhesion, cell survival and death; a process significantly involved in cancer initiation and metastasis [68, 69]. Once RHOGDI is inhibited, it induces constitutive activation of Rho GTPases, resulting in several malignant phenotype including tumour growth, angiogenesis, and invasive phenotypes [69, 70]. For instance, knocking out one of the three RHODGI genes resulted in a renal defect that progressively leads to death in adult mice, although embryonic development was not affected [71]. Togawa, Miyoshi [72] also found that male mice lacking RHOGDI1 were infertile with impaired spermatogenesis. The authors also reported problems of implantation in female mice due to this knockout. The knockout of two of the three RHOGDIs often results in a more severe phenotypes with additional immunological defects than when one of the RHOGDIs is disrupted [73]. Numerous studies have also reported that the RHOGDIs protein are involved in sperm movement, sperm capacitation and acrosome reaction, a process that is critical to occur for the sperm to interact and penetrate the egg for fertilization to take place [74-76]. Perhaps, this could be the major reason why signaling by Rho family GTPases were enriched in our metabolic pathway analysis. Notably, low reproduction performance is one of the major challenges facing beef producers in Northern Australia [77, 78]. Reproductive wastage is usually common, which is often a result of pregnancy failure and calf mortality [79, 80]. Given the role of the RHODGI pathway in reproduction, future studies could use gene expression data to investigate the genes involved in these pathways as a candidate region for infertility in cattle since we only use the **r** estimates in this study.

## Conclusion

Overall, the framework developed in this study extends our knowledge about the regions driving and antagonizing correlations between quantitative traits. About 40% of the total genomic regions were identified as driving and antagonizing genetic correlations between male and female fertility traits in the two population. These regions confirmed the polygenic nature of the traits being studied and pointed to genes of interest. Quantitative trait loci (QTL) and functional enrichment analysis revealed that many significant windows co-located with known QTLs related to milk production and fertility traits, especially puberty. In general, the enriched reproductive QTLs driving the genetic correlations between male and female fertility are the same for both cattle populations, while the antagonizing regions were population specific. Moreover, most of the antagonizing regions were mapped to the chromosome X. These results suggest regions of the chromosome X for further investigation into the trade-offs between male and female fertility. Although the methodology was applied to cattle phenotypes, using high-density SNP genotypes, the general framework developed can be applied to any species or traits, and it can easily accommodate genome sequence data.

## Materials and Methods

### Traits, genotypes and estimated genetic correlations

The traits used to demonstrate this methodology are a subset of traits from our previous study [26], where bivariate genetic correlations were estimated between 7 male and 6 female early-in-life reproductive phenotypes in two independent tropical beef cattle populations (BB and TC). The two female traits selected for this study are age at detection of the first *corpus luteum* (AGECL, days) and cows’ blood concentration of insulin growth-factor 1, measured at 18 months of age (IGF1c). Only one male trait was selected: the blood concentration of insulin growth-factor 1, measured at 6 months of age (IGF1b). These traits are important in beef cattle fertility, especially during pubertal development. The estimated genomic correlations between the traits listed above in each population have been reported in our previous study [26]. These estimates and their corresponding standard error (S.E), number of SNPs and number of animals in each population are provided in Table 2. These traits were selected because they had significant estimates of genomic correlation (i.e., traits with standard error (S.E) less than half of the size of the correlation) and different strength or direction of genetic relationships (i.e., negatively, and positively correlated traits). In brief, across-sex genetic correlations were estimated in a bivariate analysis using the linear mixed model approach. Firstly, the 770,000 genotypes were mapped to the new assembly of the bovine reference genome (ARS_UCD1.2, GenBank assembly accession GCA_002263795.2; [81]). After quality control filtering (i.e., excluding all SNPs with a minor allele frequency less than 5%), 554,712 and 686,626 SNPs remained for BB and TC datasets, respectively. Finally, bivariate genetic correlations were estimated using GIBBS2F90 [82], resulting to the estimates in Table 2.

### Overview of methods

For each trait considered in the two beef cattle populations, we estimated the genomic breeding values (GEBVs) of individuals using the genomic best linear unbiased prediction (GBLUP) model:

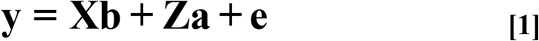

where ***y*** is the vector of phenotypes, **X** is the incidence matrix of fixed effects, ***b*** is the vector of fixed effects, **Z** is the design matrix assigned to genomic breeding values, ***a*** is the vector of GEBVs for each animal, and ***e*** is the vector of residuals. Vectors **a** and **e** are assumed to follow a normal distribution, thus **a** ∼ N (**0, Gσ**^**2**^**g** □) and **e**∼N (**0, Iσ**^**2**^**e**) □. Matrix **G** is the genomic relationship matrix.

Following the estimation of GEBVs for each animal in each trait, we then back-solved these GEBVs to obtain SNP effects for all chromosomes following the method illustrated by Strandén and Garrick [83];

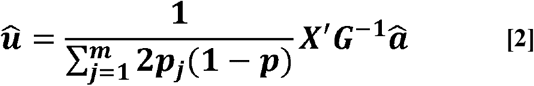

Where ***û*** is the vector of estimated SNP effects; ***m*** is the number of SNPs; ***p***_*j*_ is the allele frequency of the second allele of the jth marker; **X** is a matrix with gene contents for all markers; **G** is a genomic relationship matrix and ***â*** is a vector of GEBVs.

Notably, the estimation of GEBVs using GBLUP model [84-86] and the back-solving for GEBVs to obtain SNP effects using equation **[2]** both fits all SNP simultaneously, thereby allowing the joint estimation of SNP effects. As a result, the SNP effects from this approach are not biased by LD and resulting effect sizes can be considered independent.

Using a chromosome-wide approach, we divided SNPs on the same chromosome into small sliding windows of 500 SNPs each and then estimated the correlation (**r**) between traits as being the correlation between the 500-SNP effects estimated for trait A and the 500-SNP effects estimated for trait B. We then moved 100 SNPs further from the start of the previous window to select the next 500-SNP window, which partially overlapped with previous window, hence producing sliding windows that were 100 SNPs distant from the previous window. This was repeated for each trait pair, and for each chromosome, in a chromosome-by-chromosome approach. The resulting **r** estimates for all the chromosomes combined were denoted as W_1_….W_n_. The graphical illustration of this framework is presented in Figure 7. Moreover, the coordinates of the windows (W_1_…W_n_) were mapped to the ARS_UCD1.2 bovine reference genome. The signals across the genome were visualized with the **r** estimates of each window on the y-axis and genomic position (i.e the midpoint of the start and end position of each window) of each chromosome on the x-axis. The mapping to the bovine reference genome and plotting of the windows signals’ graphs were done using SNP & Variation Suite v8.x Golden Helix [87].

**Figure 7.**
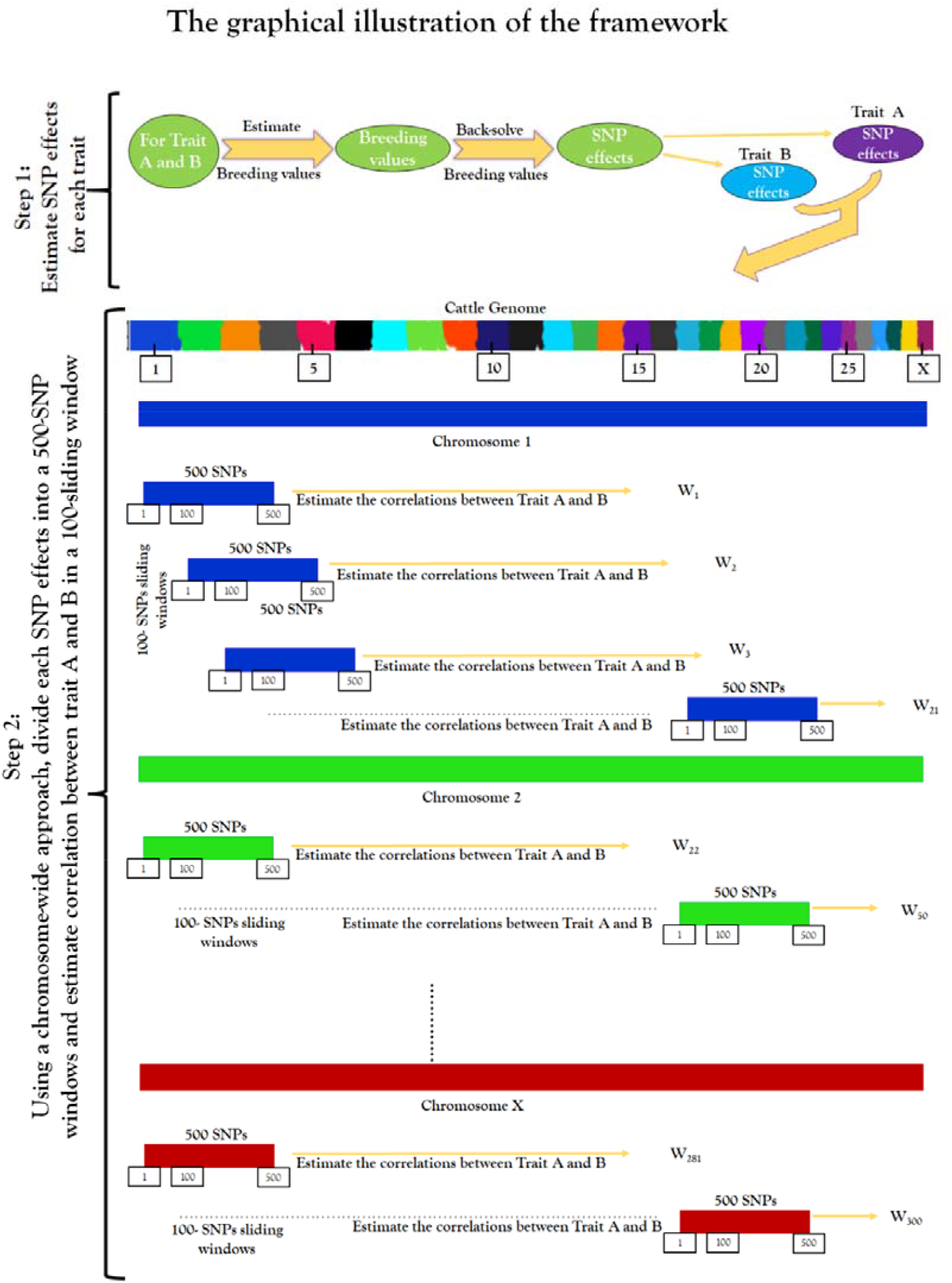
The graphical illustration of the sliding window framework. The framework involves 2 steps. Step 1 start from the estimation of genomic breeding values to the obtainment of SNP effects for each pairwise trait. Step 2 start from the estimation of 500-SNP effects in a chromosome-wide approach to obtainment of the correlation estimate in a 100-sliding window.

Depending on the overall genetic correlation observed between the traits considered, the driver and antagonizing windows can be deduced. In this study, AGECL and IGF1b were negatively correlated. Hence, the driver windows were windows with significant and negative **r** estimates, while the antagonizings were windows with significant positive **r** estimates. For the positively correlated relationship between IGF1c and IGF1b, the driver windows were windows with significant and positive **r** estimates and the antagonizings were windows with significant and negative **r** estimates. The significance of each window was established with a permutation test, described in the next section.

### Permutation test

To ensure the **r** estimates are not just noise but real signals, we performed permutation test by randomly reshuffling the SNP effects in each window across all the chromosomes in 1,000 iterations for each trait. Subsequently, we estimated correlations for 100-sliding windows of 500-SNP effects as described above. Finally, we observed the maximum and minimum **r** estimates for all the windows (W_1_…….W_n_) across the 1,000 iterations to reveal windows that were significant on the pairwise traits under investigation. Afterwards, we mapped the resulting **r** estimates for each window to the ARS_UCD1.2 bovine reference genome and plot the **r** estimates on the y-axis against the genomic position of each chromosome on the x-axis as described above. Consequently, significant windows were selected for the drivers and antagonizings genomic regions for each pairwise. Windows that were not significant were tagged “neutral windows” i.e., windows with no effect on the pairwise trait. Apart from using these windows to estimate genomic correlations and investigate the proportion of variance captured by these regions, they were excluded from other subsequent analyses. Finally, the **r** estimates of the significant windows for the driver and antagonizing regions were ranked from top to bottom in percentage (%) and the rank values were used solely for the purpose of subsequent downstream analyses. The ranking was done separately for the driver and antagonizing windows for each pairwise trait investigated in each population.

### Gene and Quantitative Traits Loci (QTL) annotation

The significant windows along with their corresponding chromosome coordinates, **r** estimates and rank values for the driver and antagonizing regions that passed the specified threshold criteria following the permutation test in BB and TC were selected. The selected windows were used for gene and QTL annotation using R package GALLO: Genomic Annotation in Livestock for positional candidate Loci (https://CRAN.R-project.org/package=GALLO) [27]. The .gtf annotation file corresponding to the bovine gene annotation from ARS-UCD1.2 assembly and the .gff file with the QTL information from cattle QTL Database (https://www.animalgenome.org/cgi-bin/QTLdb/index; [88, 89]), were used for gene and QTL annotation, respectively [27]. The two files use the same bovine reference genome (ARS-UCD1.2) to map the gene and QTLs. A remarkable advantage of GALLO is that the software retains all the information present in the input file when producing the output file. As a result, genes within each window can retain their **r** estimates and the rank values specific for their window.

The number and percentage of genes and QTLs annotated within a population (BB or TC) and the overlaps across populations (BB and TC) were investigated. Furthermore, we examined the QTLs representativeness and diversity to explain better the genomic content of the significant windows for the driver and antagonizing regions. Hence, the visualization of the percentage of cattle QTL types from cattle QTL database (i.e milk, reproduction, production, meat and carcass, health and exterior) were plotted using a pie chart by GALLO (27).

### QTL enrichment analysis

To further test the significance of the QTLs, a within population QTL enrichment analysis was conducted using a chromosome-based approach. The QTL enrichment analysis, using all the QTL information annotated within the significant windows for the driver and antagonizing regions, was performed using the qtl_enrich function from GALLO [90, 91]. Briefly, the observed number of QTLs for each trait in each annotated chromosome were compared with the expected number using a hypergeometric test approach in a 1,000 iteration rounds of random sampling from the entire cattle QTL database. With this approach, a p-value for the QTL enrichment status of each annotated QTLs within the significant windows was estimated. These estimated p-values were corrected for multiple testing using a false discovery rate (FDR) of 5%. In addition, we used chord plots to reveal the relationships between the two breeds for the enriched reproductive QTLs based on the driver and antagonizing genomic regions.

### Functional enrichment analysis

The annotated genes along with their corresponding **r** estimates and rank values for the significant driver and antagonizing windows for each pairwise trait in BB and TC populations were subjected to enrichment analysis using the commercial QIAGEN’s Ingenuity Pathway Analysis (IPA; v.8.8, http://www.ingenuity.com). The IPA allows identifying overrepresented biological mechanism, metabolic pathways, and diseases and biological functions that are highly relevant to the traits of interest using the directionality of the submitted gene list [92, 93]. The outcome of our methodology indicates that genes within each window come with their directionalities, in this case, r estimates. Thus, we leveraged on the directionality of each gene by allowing the driver genes to be upregulated and antagonizing genes to be downregulated.

Summarily, a merged dataset containing gene identifiers that were significant for both the driver and antagonizing windows for each pairwise trait in each population and their corresponding **r** estimates and rank values were uploaded into IPA. The **r** estimates were used as the “Expr Log Ratio” and the rank values were used as p-values. The IPA software recognizes gene with positive signs (+) for “Expr Log Ratio” as upregulated genes and negative sign (-) as downregulated genes. We aim to allow the driver gene lists to have positive values for “Expr Log Ratio” and the antagonizing gene lists to be negative. Where this is not achievable based on the original **r** estimates (i.e., AGECL vs IGF1b), we reversed the sign for the driver and antagonizing genes to meet this objective.

Of note, IPA can only analyse a maximum of 8,000 gene list. In most cases, the merged gene list for each pairwise trait in each population is often >8,000. Hence, we used the rank values as the cut-off to select the top ∼80% genes from the driver and antagonizing gene list for the pathway analyses. Using a proportion of the gene list to infer biological pathways might result in the loss of some important biological information relevant to the trait of interest. We analysed the driver and antagonizing gene list separately for each pairwise trait in each population to ensure no important information was lost because of the cut-offs. Further, we compare the result of the separate analyses with the merged gene list from the ∼80% cut-off.

The pathway analysis was conducted using the “Core Analysis” function implemented within IPA. In this analysis, associations were calculated using direct and indirect relationships among the gene lists. At first, the gene lists were mapped to human gene data. Genes without an associated gene symbol or gene annotation were subjected to an annotation by homology using BioMart application available in the Ensembl database (http://www.ensembl.org/biomart/martview/) [94, 95]. With this approach, we only considered non-annotated genes with percentage of identity ≥80% with human homolog. The final datasets used for the IPA analyses are presented in Additional file 8 (Table S31-34). Finally, the “Core Analysis” was used to identify canonical metabolic pathways enriched at Benjamini–Hochberg corrected p-values (B-H-P-value) of p<0.01).

## Supporting information

Additional file 1: The number of windows, chromosome number, chromosome coordinates, and correlation estimates for each window for the two pairwise trait in Brahman and Tropical Composite population (Table S1-S2).

Additional file 2: Genome plots of the correlation estimates from the permutation test at 500 iterations in BB and TC for the two pairwise traits. The correlation estimates were plotted on the y-axis and the genomic position of each chromosome on the x-axis, according to the ARS_UCD1.2 bovine reference genome (Figure S1).

Additional file 3: The number of windows, chromosome number, chromosome coordinates, and correlation estimates as well as the maximum and minimum correlation estimate for each window for all trait pairwise in Brahman and Tropical Composite population following permutation test of 500-SNP effects in 100-SNP sliding windows at 1000 iterations (Table S3-S6).

Additional file 4: The number of windows, chromosome number, chromosome coordinates, correlation estimates and rank value for driver, antagonizing and the neutral regions that passed the threshold after permutation test in Brahman (BB) and Tropical Composite population for the studied trait (Table S7-18).

Additional file 5: The enriched QTLs of the driver and antagonizing regions for all trait pairwise in Brahman and Tropical Composite cattle. The enriched QTLs are rank based on the adj.pval (Table S19-26).

Additional file 6: Canonical pathways significantly enriched for all trait pairwise in Brahman and Tropical Composite population. Significantly enriched canonical pathways were identified using Benjamini-Hochberg p-values <0.01 (Figure S2-S9).

Additional file 7: The list of the significant enriched canonical metabolic pathways showing all the genes involved in each pathway for all trait pairwise in Brahman and Tropical Composite population. Significantly enriched canonical pathways were identified using Benjamini-Hochberg p-values <0.01. Z-score >2 denote the activation of the pathway. Z-score <-2 indicate the inhibition of the pathway (Table S27-30).

Additional file 8: The final dataset used for Ingenuity Pathway Analysis (IPA) for all trait pairwise in Brahman and Tropical Composite cattle (Table S31-34).

## Abbreviations

AGECL: Age at first corpus luteum
BB: Brahman
BTA: Bovine chromosome
GBLUP: Genomic Best Linear Unbiased Prediction
GEBV: Genomic estimated breeding values
IGF1b: Serum levels of insulin growth hormone measured in bulls at 6-months of age
IGF1c: Serum levels of insulin growth hormone measured in bulls at 18-months of age
IPA: Ingenuity Pathway Analysis
LD: Linkage Disequilibrium
QTL: Quantitative Trait Loci
R: Correlation estimate
SNP: Single Nucleotide Polymorphism
TC: Tropical Composite

## Declarations

### Ethics approval and consent to participate

Not applicable

### Consent for publication

Not applicable

### Availability of data and materials

The data used in this are available from the Cooperative Research Centre for Beef Genetic Technologies (Beef CRC). Data are available from https://www.beefcrc.com with the permission of Meat and Livestock Australia and the University of Queensland (if interested please contact the corresponding author).

### Authors’ contributions

O.B.S and L.P.N. conceived the idea. O.B.S wrote the code, conducted the analyses, and wrote the manuscript. M.S.T. and G.C.C. contributed to the analyses, manuscript writing and editing. A.C. conceived the downstream biological analyses. L.R., B.J.H., and M.R.S. supervised the study. All authors read, edited, and approved the final manuscript.

## Acknowledgements

This research was conducted using the legacy database of the Cooperative Centre for Beef Genetics Technologies and their core partners including Meat and Livestock Australia. The authors acknowledge all the participants of the Cooperative Centre for Beef Genetics Technologies for their efforts in conducting the field trials and recording the phenotypes. We also acknowledge the mentorship of Professor Steve Moore to Babatunde Olasege in his PhD.

