## Additional file 2 for "Correlation scan: identifying genomic regions that affect genetic correlations applied to fertility traits"

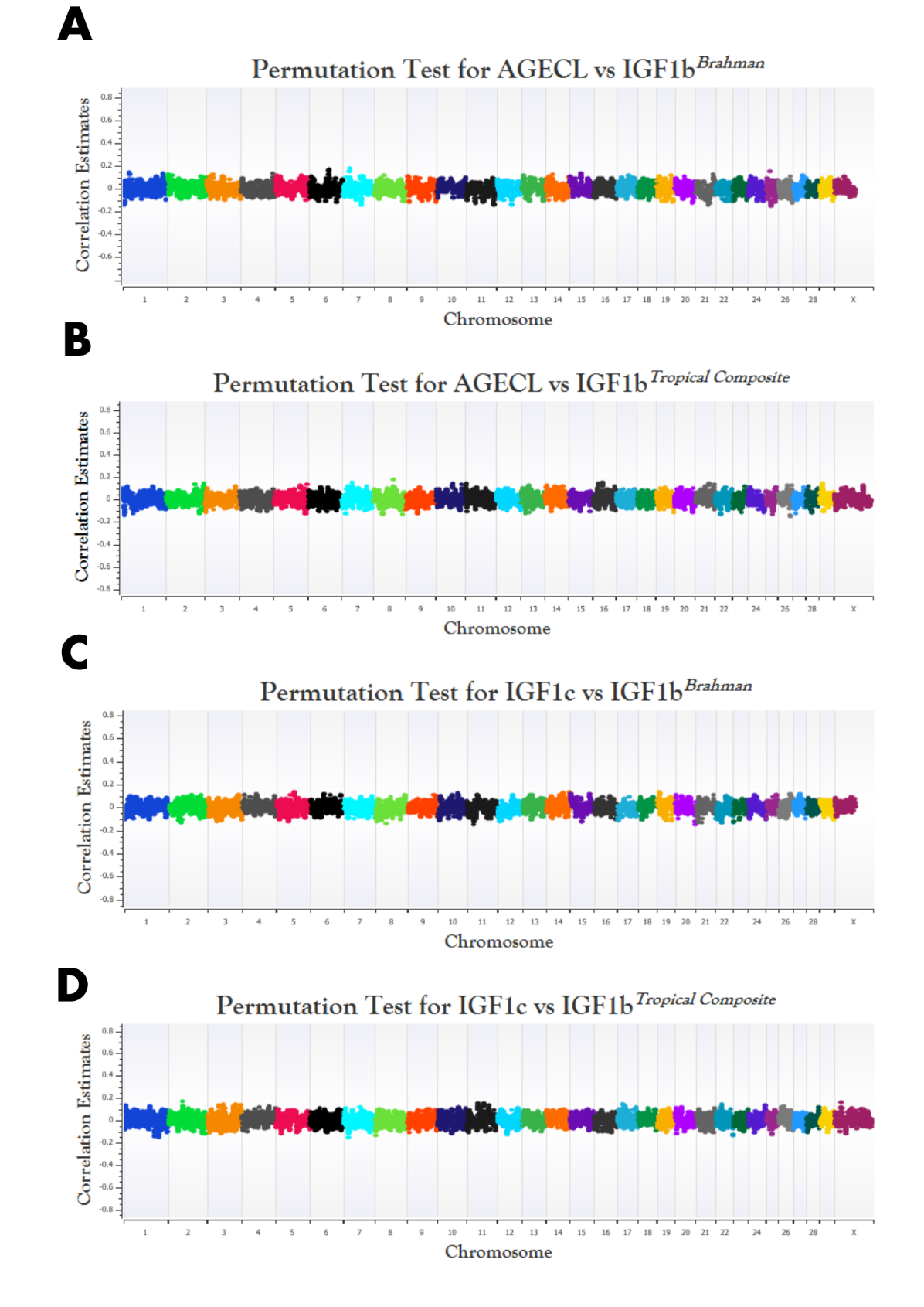

**Figure S1.** Genome plots of the correlation estimates from the permutation test at 500 iterations in BB and TC for the two pairwise traits (BB-AGECL vs IGF1b; **A**, TC-AGECL vs IGF1b; **B**, BB-IGF1c vs IGF1b; **C**, TC- IGF1c vs IGF1b; **D**). **AGECL**, age at first corpus; **IGF1**, serum levels of insulin growth hormone (measured in bulls, **IGF1b**, or cows, **IGF1c**). The correlation estimates were plotted on the y-axis and the genomic position (i.e., midpoint between the start and end position of each window) of each chromosome on the x-axis, according to the ARS_UCD1.2 bovine reference genome.
