## Additional file 6 for "Correlation scan: identifying genomic regions that affect genetic correlations applied to fertility traits"

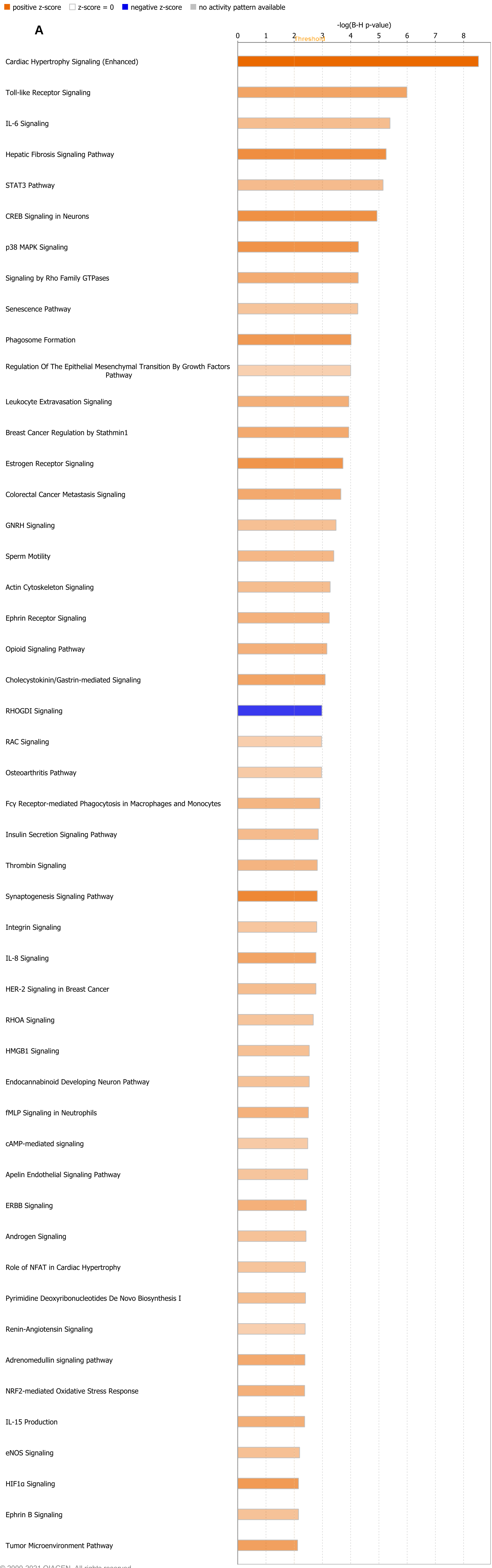


© 2000-2021 QIAGEN. All rights reserved.

**Figure S2.** Canonical pathways significantly enriched for AGECL vs IGF1b in Brahman population. Significantly enriched canonical pathways were identified using Benjamini-Hochberg pvalues <0.01. AGECL, age at first corpus; IGF1b, serum levels of insulin growth hormone measured in bulls).The left hand side is the enriched canonical pathway and the right hand side is the predicted activation state of the canonical pathway using a z-score. Orange color indicate a positive z-score (Z>2) denoting the activation of the pathway. Blue color indicate a negative z-score (Z<-2) indicating the inhibition of the pathway.

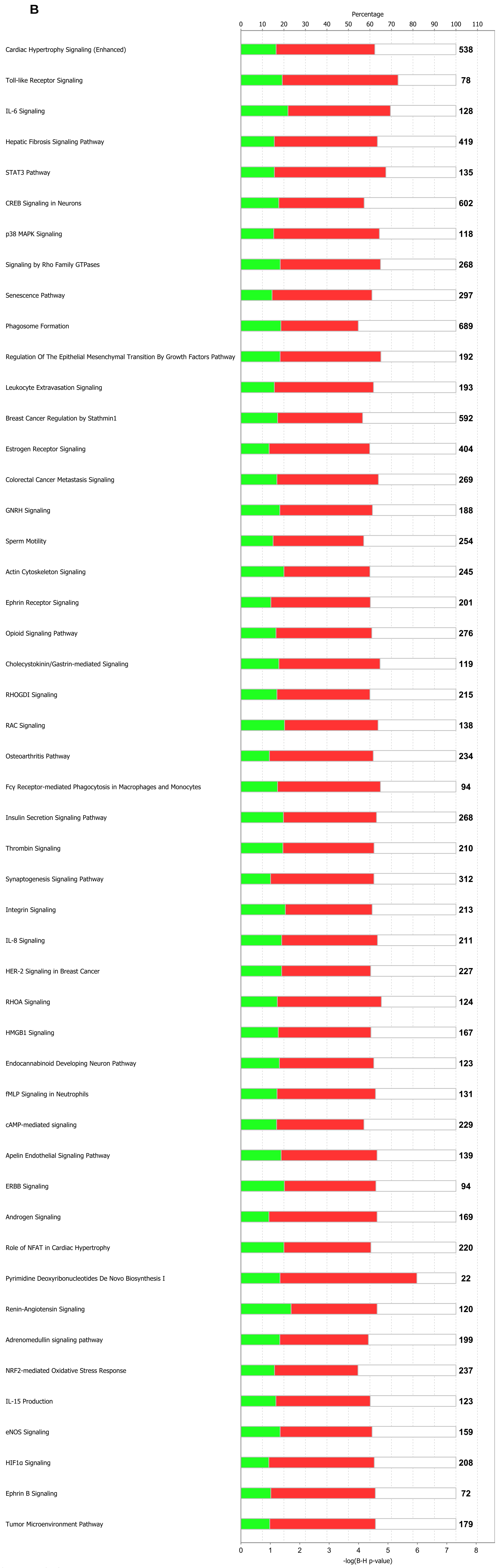


© 2000-2021 QIAGEN. All rights reserved.

**Figure S3.** The significantly enriched canonical pathways showing the percentage of driver (red) and antagonizing (green) genes in each pathway for AGECL vs IGF1b in Brahman population. Significantly enriched canonical pathways were identified using Benjamini-Hochberg pvalues <0.01. AGECL, age at first corpus; IGF1b, serum levels of insulin growth hormone measured in bulls).The left hand side is the enriched canonical pathway and the right hand side is the percentage of the driver and the antagonizing genes in the pathway

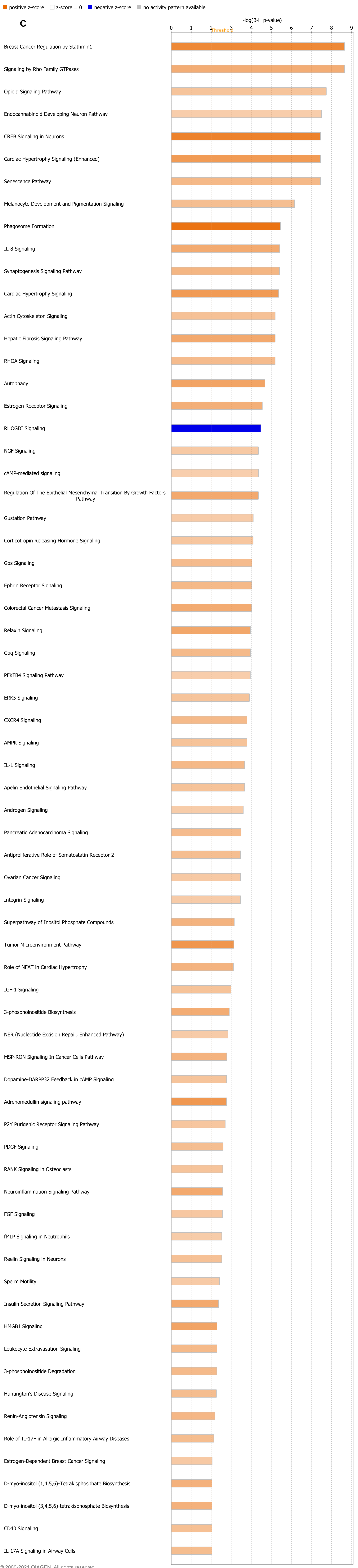


© 2000-2021 QIAGEN. All rights reserved.

**Figure S4.** Canonical pathways significantly enriched for AGECL vs IGF1b in Tropical Composite population. Significantly enriched canonical pathways were identified using Benjamini-Hochberg pvalues <0.01. AGECL, age at first corpus; IGF1b, serum levels of insulin growth hormone measured in bulls).The left hand side is the enriched canonical pathway and the right hand side is the predicted activation state of the canonical pathway using a z-score. Orange color indicate a positive z-score (Z>2) denoting the activation of the pathway. Blue color indicate a negative z-score (Z<-2) indicating the inhibition of the pathway.

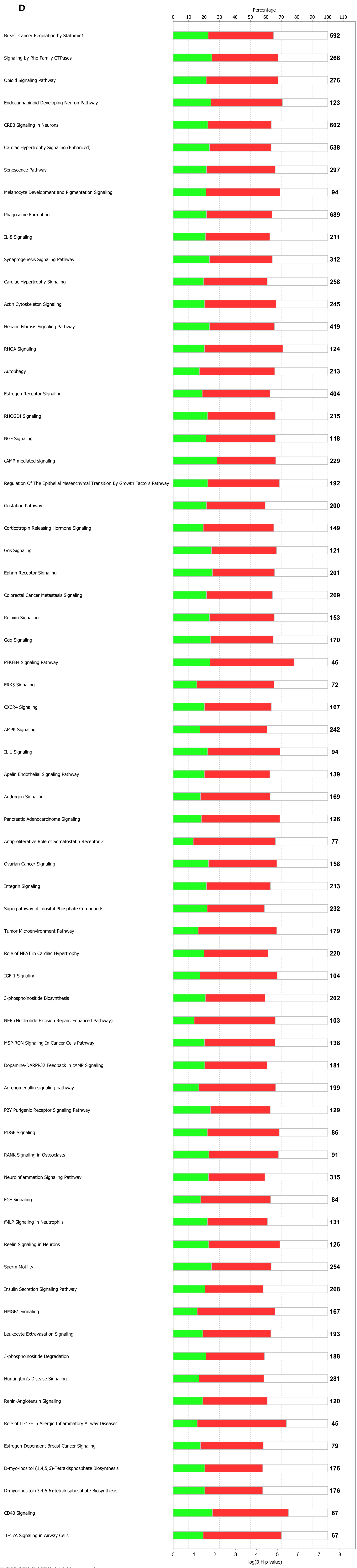


© 2000-2021 QIAGEN. All rights reserved.

**Figure S5.** The significantly enriched canonical pathways showing the percentage of driver (red) and antagonizing (green) genes in each pathway for AGECL vs IGF1b in Tropical Composite population. Significantly enriched canonical pathways were identified using Benjamini-Hochberg pvalues <0.01. AGECL, age at first corpus; IGF1b, serum levels of insulin growth hormone measured in bulls).The left hand side is the enriched canonical pathway and the right hand side is the percentage of the driver and the antagonizing genes in the pathway

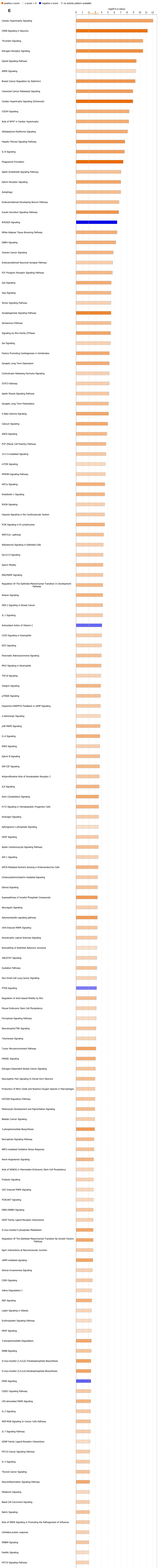


© 2000-2021 QIAGEN. All rights reserved.

**Figure S6.** Canonical pathways significantly enriched for IGF1c vs IGF1b in Brahman population. Significantly enriched canonical pathways were identified using Benjamini-Hochberg pvalues <0.01. IGF1, serum levels of insulin growth hormone (measured in bulls, IGF1b, or cows, IGF1c).The left hand side is the enriched canonical pathway and the right hand side is the predicted activation state of the canonical pathway using a z-score. Orange color indicate a positive z-score (Z>2) denoting the activation of the pathway. Blue color indicate a negative z-score (Z<-2) indicating the inhibition of the pathway.

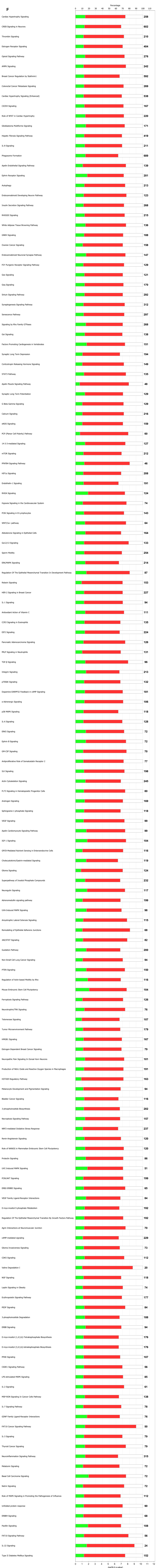


© 2000-2021 QIAGEN. All rights reserved.

**Figure S7** The significantly enriched canonical pathways showing the percentage of driver (red) and antagonizing (green) genes in each pathway for IGF1c vs IGF1b in Brahman population. Significantly enriched canonical pathways were identified using Benjamini-Hochberg pvalues <0.01. IGF1, serum levels of insulin growth hormone (measured in bulls, IGF1b, or cows, IGF1c).The left hand side is the enriched canonical pathway and the right hand side is the percentage of the driver and the antagonizing genes in the pathway

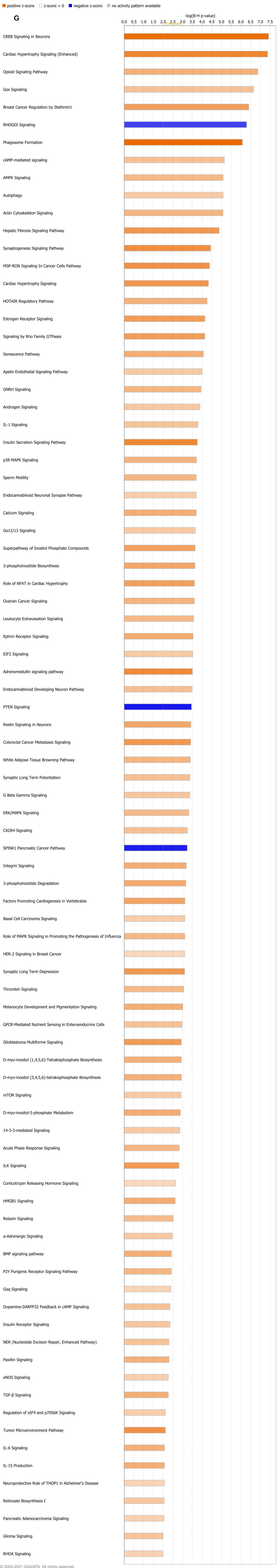


© 2000-2021 QIAGEN. All rights reserved.

**Figure S8.** Canonical pathways significantly enriched for IGF1c vs IGF1b in Tropical Composite population. Significantly enriched canonical pathways were identified using Benjamini-Hochberg pvalues <0.01. IGF1, serum levels of insulin growth hormone (measured in bulls, IGF1b, or cows, IGF1c).The left hand side is the enriched canonical pathway and the right hand side is the predicted activation state of the canonical pathway using a z-score. Orange color indicate a positive z-score (Z>2) denoting the activation of the pathway. Blue color indicate a negative z-score (Z<-2) indicating the inhibition of the pathway.

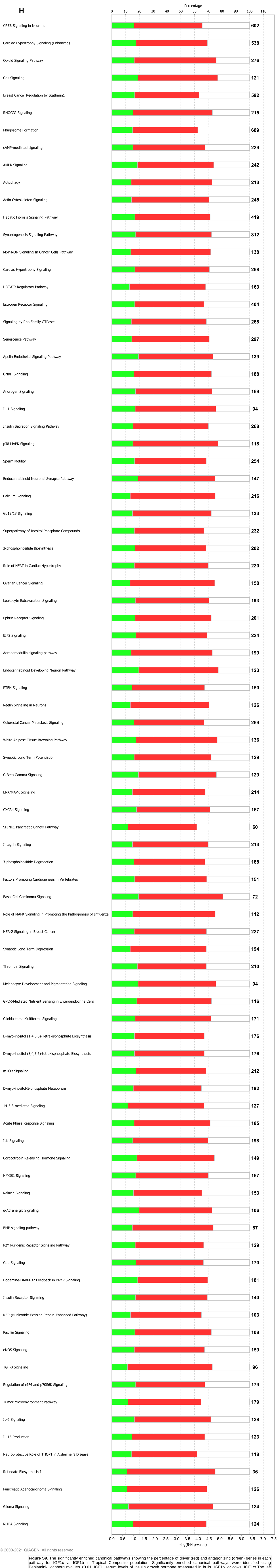


© 2000-2021 QIAGEN. All rights reserved.

**Figure S9.** The significantly enriched canonical pathways showing the percentage of driver (red) and antagonizing (green) genes in each pathway for IGF1c vs IGF1b in Tropical Composite population. Significantly enriched canonical pathways were identified using Benjamini-Hochberg pvalues <0.01. IGF1, serum levels of insulin growth hormone (measured in bulls, IGF1b, or cows, IGF1c).The left hand side is the enriched canonical pathway and the right hand side is the percentage of the driver and the antagonizing genes in the pathway
